# Evaluation of DNA extraction protocols from liquid-based cytology specimens for studying cervical microbiota

**DOI:** 10.1101/2020.01.27.921619

**Authors:** Takeo Shibata, Mayumi Nakagawa, Hannah N. Coleman, Sarah M. Owens, William W. Greenfield, Toshiyuki Sasagawa, Michael S. Robeson

**Author notes:** Corresponding author: Michael S. Robeson II (M.S.R.).

## Abstract

Cervical microbiota (CM) are considered an important factor affecting the progression of cervical intraepithelial neoplasia (CIN) and are implicated in the persistence of human papillomavirus (HPV). Collection of liquid-based cytology (LBC) samples is routine for cervical cancer screening and HPV genotyping and can be used for long-term cytological biobanking. We sought to determine whether it is possible to access microbial DNA from LBC specimens, and compared the performance of four different extraction protocols: (ZymoBIOMICS DNA Miniprep Kit; QIAamp PowerFecal Pro DNA Kit; QIAamp DNA Mini Kit; and IndiSpin Pathogen Kit) and their ability to capture the diversity of CM from LBC specimens. LBC specimens from 20 patients (stored for 716 ± 105 days) with CIN values of 2 or 3 were each aliquoted for each of the four kits. Loss of microbial diversity due to long-term LBC storage could not be assessed due to lack of fresh LBC samples. Comparisons with other types of cervical sampling were not performed. We observed that all DNA extraction kits provided equivalent accessibility to the cervical microbial DNA within stored LBC samples. Approximately 80% microbial genera were shared among all DNA extraction protocols. Potential kit contaminants were observed as well. Variation between individuals was a significantly greater influence on the observed microbial composition than was the method of DNA extraction. We also observed that HPV16 was significantly associated with community types that were not dominated by *Lactobacillus iners*.

## Introduction

High-throughput sequencing (HTS) technology of 16S rRNA gene amplicon sequences has made it possible to better understand the relationships between cervicovaginal microbiota and human papillomavirus (HPV) infection [1] [2] [3] [4] [5] and HPV-related diseases [6] [7] [8] [9] [10]. Cervicovaginal microbiota are considered to be an important factor affecting the progress of cervical intraepithelial neoplasia (CIN) [6] [7] [8] [9] and are implicated in the persistence of high-risk HPV (HR-HPV) [1] [2] and low-risk HPV (LR-HPV) [3]. However, microbial signatures associated with either HR-HPV or LR HPV can vary depending on the population under study, e.g., the phyla *Actinobacteria* and *Fusobacteria* were found to be enriched in HR-HPV positive Chinese women [4] while another study observed these groups associated with low-risk HPV (LR-HPV) in South African women [3]. Additionally, *Lactobacillus iners*-dominant samples are associated with both HR-HPV and LR-HPV [5], often associated with moderate CIN risk [10]. Moreover, it has been shown that CIN risk was increased in patients with HR-HPV [10] when the cervical microbes *Atopobium vaginae*, *Gardnerella vaginalis*, and *Lactobacillus iners* were present in greater proportion compared to *L. crispatus*. The cervicovaginal microbiome is often described by the abundance of *Lactobacillus spp., i.e.* the community is either referred to as a *Lactobacillus*-dominant type or non-*Lactobacillus*-dominant type, and can interact with the immune system in different ways [7] [11]. For example, inflammatory cytokines, such as Interleukin (IL)-1α and IL-18, were increased in non-*Lactobacillus*-dominant community types of reproductive-aged healthy women [11]. In the analysis of patients with cervical cancer, non-*Lactobacillus*-dominant community types were positively associated with chemokines such as interferon gamma-induced protein 10 (IP-10) and soluble CD40-ligand activating dendric cells (DCs) [7]. The metabolism of the cervicovaginal microbiome may be a substantial contributing factor to maternal health during pregnancy, although the mechanism is still unclear [12]. Prior research, on the importance of the microbiome in cancer therapeutics via checkpoint inhibitors [13], along with our own work on the role of CM in vaccine response [14], suggests that the CM has a significant role to play in disease progression and therapeutic treatment. We continue our work here to further assess use liquid-based cytology (LBC) samples to survey microbial community DNA.

Little has been reported on the utility of LBC samples for use in cervical microbiome studies. Conventionally, microbiome sample collection methods entail the use of swabs [15] or self-collection of vaginal discharge [16]. To obtain a non-biased and broad range of cervical microbiota, DNA extraction should be optimized for a range of difficult-to-lyse-bacteria, *e.g*. *Firmicutes*, *Actinobacteria*, and *Lactobacillus* [15] [17] [18] [19] [20].

LBC samples are promising for cervicovaginal microbiome surveys, as they are an already established method of long-term cytological biobanking [21]. In clinical practice, cervical cytology for cervical cancer screening or HPV genotyping is widely performed using a combination of cervical cytobrushes and LBC samples such as ThinPrep (HOLOGIC) or SurePath (BD). An LBC specimen can be used for not only cytological diagnosis but also additional diagnostic tests such as HPV, *Chlamydia*, *Neisseria gonorrhoeae*, and *Trichomonas* infection [22] [23] [24]. Despite the promising potential to use LBC samples for surveying cervicovaginal microbiota, it is known that DNA contained within LBC samples may degrade over prolonged storage times when kept at ambient or non-freezing temperatures [25] [26]. Although others have shown minimal DNA degradation of LBC samples stored at −80□C [26], the ability to reliably access microbial DNA remains to be seen, and is the focus of our study.

Furthermore, the ability to characterize these microbiota, as commonly assessed by 16S rRNA gene sequencing, can be biased as a result of methodological differences of cell lysis and DNA extraction protocols [27] [28] [29]. Herein, we compare four different commercially available DNA extraction kits in an effort to assess their ability to extract and characterize any viable microbial DNA from stored LBC samples. Additionally, we examine the relationship between HPV infection and the composition of cervical microbiota still accessible after prolonged LBC storage.

## Methods

### Recruiting patients

Patients were participants of a single center randomized double blind Phase II clinical trial (NCT02481414) in which enrollees were assigned to receive an HPV therapeutic vaccine called PepCan or an adjuvant derived from *Candida albicans* (Candin®, Nielsen BioSciences, San Diego, CA). Pre-injection liquid based cervical cytology (ThinPrep) samples from 20 consecutive enrollees who gave written informed consent between 3/21/2017 and 12/11/2017 were used for this study. Patients were recruited mainly through referrals from clinics from inside and outside of the medical center. Flyers and Google advertisements were also utilized. Inclusion Criteria: aged 18–50 years, had recent (≤ 60 days) Pap smear result consistent with HSIL or “cannot rule out HSIL” or HSIL on colposcopy guided biopsy, untreated for HSIL or “Cannot rule out HSIL”, able to provide informed consent, willingness and able to comply with the requirements of the protocol. Exclusion Criteria: history of disease or treatment causing immunosuppression (e.g., cancer, HIV, organ transplant, autoimmune disease), being pregnant or attempting to be pregnant within the period of study participation, breast feeding or planning to breast feed within the period of study participation, allergy to Candida antigen, history of severe asthma requiring emergency room visit or hospitalization within the past 5 years, history of invasive squamous cell carcinoma of the cervix, history of having received PepCan. Those who qualified for the study based on their cervical cytology underwent cervical biopsy, and they qualified for vaccination if the results were CIN2/3. All collected samples are representative of a larger population in gynecology clinics with abnormal Pap tests. If in the opinion of the Principal Investigator or other Investigators, it is not in the best interest of the patient to enter this study, the patient was excluded. Patients’ age, race, and ethnicity were recorded based on standard NIH requirements. All categories and definitions, *e.g.* ethnicity, age, etc., were based on NIH Guidelines.

### Sampling of cervical microbiome

The cervical cytology specimens in this current study were collected before the vaccination and reserved in the vial of the ThinPrep Pap Test (HOLOGIC) as described in Ravilla *et al*. 2019 [14]. The specimens were frozen in an ultra-low temperature freezer (−80□C) on the day of collection. The storage period from sample collection to DNA extraction was 716 ± 105 days.

### HPV genotyping

HPV-DNA was detected by Linear Array HPV Genotyping Test (Roche Diagnostics) which can detect up to 37 HPV genotypes using ThinPrep solution [30]. HPV16, 18, 31, 33, 35, 39, 45, 51, 52, 56, 58, 59, and 68 were defined as HR-HPV genotypes; and HPV6, 11, 40, 42, 54, 61, 62, 71, 72, 81, 83, 84, and CP6108 were defined as LR-HPV genotypes [31] [32].

### DNA extraction protocols

We selected four commercially available DNA extraction kits as the candidates for comparison: ZymoBIOMICS DNA Miniprep Kit (Zymo Research, D4300), QIAamp PowerFecal Pro DNA Kit (QIAGEN, 51804), QIAamp DNA Mini Kit (QIAGEN, 51304), and IndiSpin Pathogen Kit (Indical Bioscience, SPS4104). These kits have been successfully used in a variety of human cervical, vaginal, and gut microbiome surveys [10] [21] [33]. We’ll subsequently refer to each of these kits in abbreviated form as follows: ZymoBIOMICS, PowerFecalPro, QIAampMini, and IndiSpin. The protocols and any modifications are outlined in Table 1.

**Table 1:**
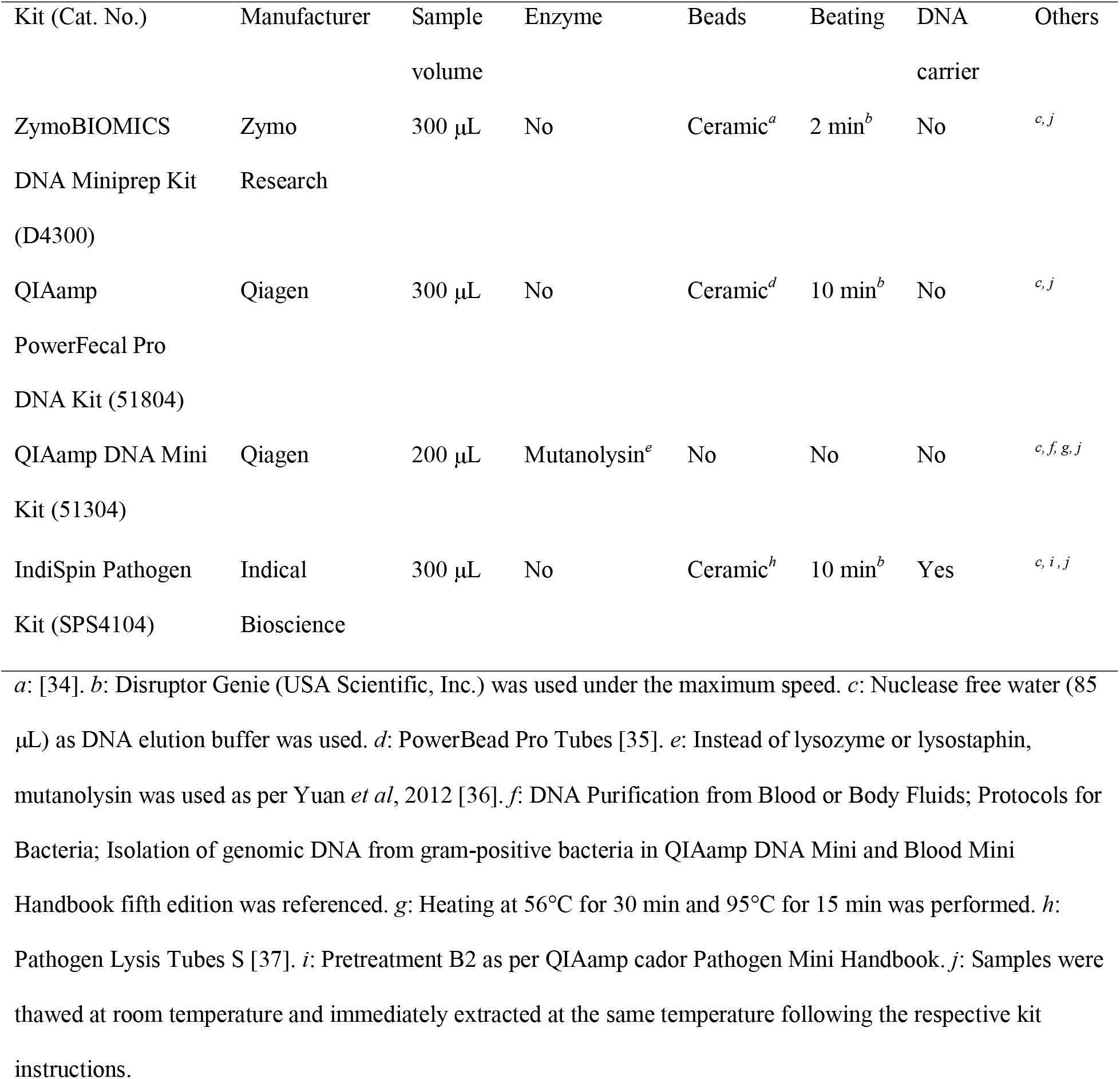
Characteristics of four different DNA extraction protocols.

Each LBC sample was dispensed into four separate 2 mL sterile collection tubes (dispensed sample volume = 500 μL) to create four cohorts of 20 DNA extractions (Figure 1). Each extraction cohort was processed through one of the four kits above. A total of 80 extractions (4 kits × 20 patients) were prepared for subsequent analyses. Applied sample volume of ThinPrep solution was 300 μL for ZymoBIOMICS, 300 μL for PowerFecalPro, 200 μL for QIAampMini, and 300 μL for IndiSpin. The sample volume was standardized to 300 μL as long as the manufacturer’s instructions allowed to do so. DNA extraction for all samples was performed by the same individual who practiced by performing multiple extractions for each kit before performing the actual DNA extraction on the samples analyzed in this study. Positive control was mock vaginal microbial communities composed of a mixture of genomic DNA from the American Type Culture Collection (ATCC MSA1007). Negative control was the ThinPrep preservation solution without the sample as blank extraction [38].

**Figure 1.**
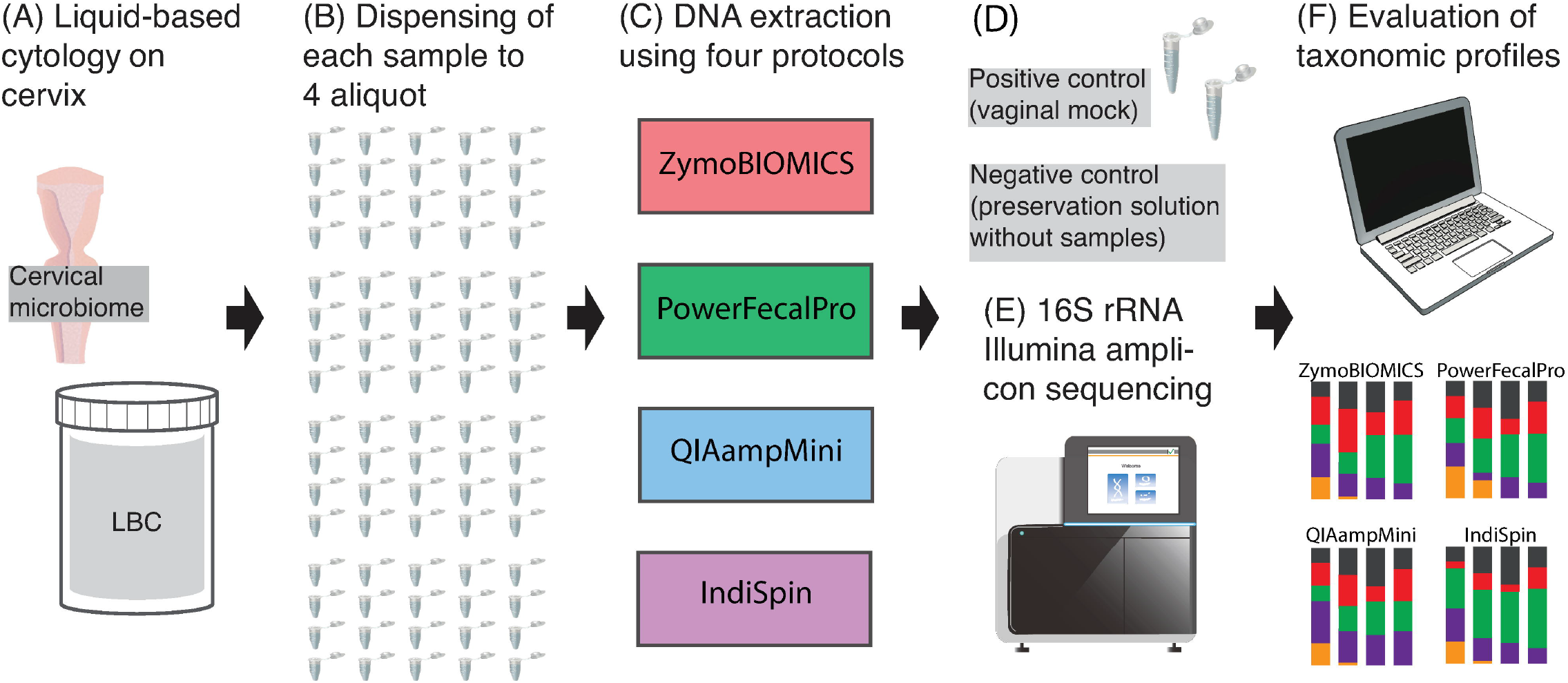
Overview of the study design using 16S rRNA gene to compare the DNA extraction protocol. (A) Liquid-based cytology (LBC) specimens from 20 patients with CIN2/3 or suspected CIN2/3. (B) A total of 80 DNA extractions were performed. (C) The four DNA extraction methods. (D) DNA of mock vaginal community as a positive control and preservation solution as a negative control. (E) Sequencing using Illumina MiSeq. (F) Analysis of the taxonomic profiles among the DNA extraction protocols. Images form Togo Picture Gallery [39] were used to create this figure.

### Measurement of DNA yield

DNA yield for each method was evaluated by spectrophotometer (Nanodrop One, Thermo Scientific). Analysis of the DNA yield from IndiSpin was omitted as nucleic acid is used as a carrier for this kit. The mean DNA yields per 100 μL ThinPrep sample volume were compared.

### 16S rRNA marker gene sequencing

Controls and the extracted DNA were sent to Argonne National Laboratory (IL, USA) for amplification and sequencing of the 16S rRNA gene on an Illumina MiSeq sequencing platform [40]. The same volume of DNA was used for each reaction, and then normalized at the PCR pooling step. This ensures that equal amounts of each amplified sample are added to the sequencing pool. Paired-end reads from libraries with ∼250-bp inserts were generated for the V4 region using the barcoded primer set: 515FB: 5’-GTGYCAGCMGCCGCGGTAA-3’ and 806RB: 5’-GGACTACNVGGGTWTCTAAT-3’ [41] [42] [43] [44] [45]. MiSeq Reagent Kit v2 (2 × 150 cycles, MS-102-2002) was used.

### Sequence processing and analysis

Initial sequence processing and analyses were performed using QIIME 2 [46], any commands prefixed by `q2-` are QIIME 2 plugins. After demultiplexing of the paired-end reads by `q2-demux`, the imported sequence data was visually inspected via QIIME 2 View [47], to determine the appropriate trimming and truncation parameters for generating Exact Sequence Variants (ESVs) [48] via `q2-dada2` [49]. ESVs will be referred to as Operational Taxonomic Units (OTUs). The forward reads were trimmed at 15 bp and truncated at 150 bp; reverse reads were trimmed at 0 bp and truncated at 150 bp. The resulting OTUs were assigned taxonomy through `q2-feature-classifier classify-sklearn`, by using a pre-trained classifier for the amplicon region of interest [50]. This enables more robust taxonomic assignment of the OTUs [51]. Taxonomy-based filtering was performed by using `q2-taxa filter-table` to remove any OTUs that were classified as “Chloroplast”, “Mitochondria”, “Eukaryota”, “Unclassified” and those that did not have at least a Phylum-level classification. We then performed additional quality filtering via `q2-quality-control`, and only retained OTUs that had at least a 90% identity and 90% query alignment to the SILVA reference set [52]. Then `q2-alignment` was used to generate a *de novo* alignment with MAFFT [53] which was subsequently masked by setting `max-gap-frequency 1 min-conservation 0.4`. Finally, `q2-phylogeny` was used to construct a midpoint-rooted phylogenetic tree using IQ-TREE [54] with automatic model selection using ModelFinder [55]. Unless specified, subsequent analyses were performed after removing OTUs with a very low frequency of less than 0.0005% of the total data set in this case [56].

### Microbiome analysis

To compare the taxonomic profiles among four types of DNA extraction methods (Figure 1 & Table 1), the following analyses were performed; (I) bacterial microbiome composition, (II) detection of common and unique taxa, (III) alpha and beta diversity analysis, and (IV) identification of specific bacteria retained per DNA extraction method.

Overall microbial composition was investigated at the family and genus taxonomic level. After all count data of taxonomy were converted to relative abundance, the top 10 abundant taxonomic groups in each family and genus level were plotted in colored bar plot [57] [58] [59]. Variation of microbiome composition per DNA extraction method or per individual was assessed by the Adonis test (`q2-diversity adonis`) [60] [61].

We set out to determine which microbial taxonomic groups were differentially accessible across the sampling protocols by LEfSe analyses [62]. We further assessed the microbial taxa using jvenn [63] at family and genus level. The Venn diagram was created after removing OTUs with a frequency of less than 0.005% [56]. We used Scheffe test [64] to perform post-hoc analysis of the LEfSe output.

Analytical approaches (at the OTU-level) that do not require the rarefying of data, such as `q2-breakaway` [65] and Aitchison distance using `q2-deicode` [66] were used to determine both alpha (within-sample) and beta (between-sample) diversity respectively. These were compared with traditional methods, that often require the data to be rarefied. Here we applied the following traditional alpha and beta-diversity metrics: Faith’s Phylogenetic Diversity, Observed OTUs, Shannon’s diversity index, Pielou’s Evenness, Unweighted UniFrac distance, Weighted UniFrac distance, Jaccard distance, and Bray-Curtis distances via `q2-diversity` [46]. In order to maintain a reasonable balance between sequencing depth and sample size, we determined that a rarefaction depth of 51,197 reads allowed us to retain data for all four kits for 15 of the 20 individual patients. Overall, our subsequence analysis consisted of 3,071,820 reads (27.6%, 3,071,820 / 11,149,582 reads). All diversity measurements used in this study are listed in Table S1.

### Community type and HPV status

In addition to the analysis above, we tested whether the samples clustered by microbiome composition were related to the patient’s clinical and demographic characteristics such as, cervical biopsy diagnosis, race, and HPV16 status. HPV16 status has been reported to be associated with both racial differences as well as microbial community types [14] [67] [68] [69]. We employed the DMM [70] model to determine the number of community types for bacterial cervical microbiome. Then, we clustered samples to the community type [9] [71]. Since vaginal microbiota were reported to be clustered with different *Lactobacillus sp*. such as *L. crispatus*, *L. gasseri*, *L. iners*, or *L. jensenii* [18] [72], we also collapsed the taxonomy to the species level and performed a clustering analysis using “microbiome” R package [59]. We then determined which bacterial taxa were differentially abundant among the patients with or without HPV16 via `q2-aldex2` [73] and LEfSe [62].

### General statistical analysis

All data are presented as means ± standard deviation (SD). Comparisons were conducted with Fisher’s exact test or Dunn’s test with Benjamini-Hochberg-adjustment [74] or Wilcoxon test with Benjamini-Hochberg-adjustment or pairwise PERMANOVA when appropriate. A p value < 0.05 or a q value < 0.05 was considered statistically significant. We did not control for confounding variables such as socioeconomic status, nutrition, environmental exposures, or similar factors.

### Ethics approval and consent to participate

This study was approved by the Institutional Review Board at the University of Arkansas for Medical Sciences (IRB # 202790). No minors were enrolled in this study.

### Consent for publication

Written informed consent for publication was obtained for all patients under IRB # 202790; NCT # NCT02481414; IND # 15173.

### Availability of data and materials

MIMARKS compliant [75] DNA sequencing data are available via the Sequence Read Archive (SRA) at the National Center for Biotechnology Information (NCBI), under the BioProject Accession: PRJNA598197.

## Results

### Patient characteristics

The age of the patients (n = 20) was 31.4 ± 5.0 years. The distribution of race was 15% African American (n = 3), 50% European descent (n = 10), and 35% Hispanic (n = 7). Cervical histology was 40% CIN2 (n = 8), 50% CIN3 (n = 10), and 10% benign (n = 2). HPV genotypes were 50% HPV16 positive (n = 10), 10% HPV18 positive (n = 2), 90% HR-HPV positives (n = 18), 45% LR-HPV positives (n = 9), and 75% multiple HPV positives (n = 15). Patient characteristics were summarized in Table 2.

**Table 2.**
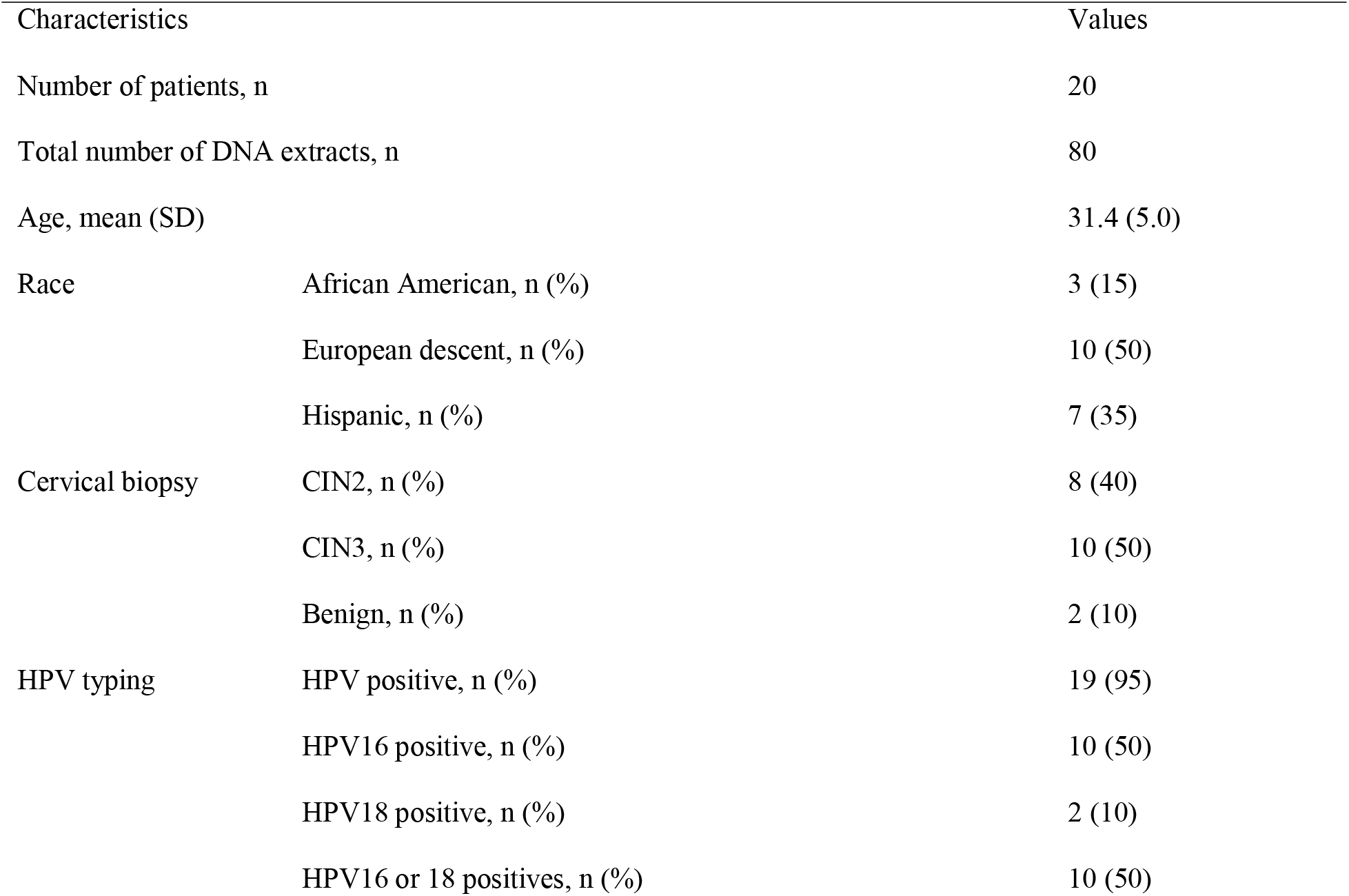
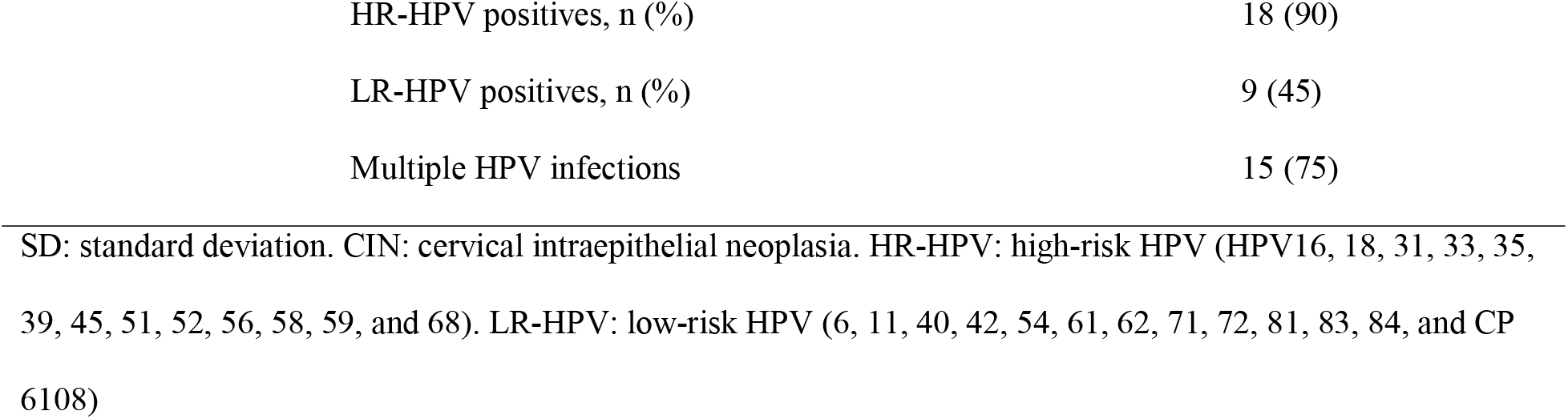
Patient characteristics.

| Characteristics |  | Values |
| --- | --- | --- |
| Number of patients, n |  | 20 |
| Total number of DNA extracts, n |  | 80 |
| Age, mean (SD) |  | 31.4 (5.0) |
| Race | African American, n (%) | 3 (15) |
|  | European descent, n (%) | 10 (50) |
|  | Hispanic, n (%) | 7 (35) |
| Cervical biopsy | CIN2, n (%) | 8 (40) |
|  | CIN3, n (%) | 10 (50) |
|  | Benign, n (%) | 2 (10) |
| HPV typing | HPV positive, n (%) | 19 (95) |
|  | HPV16 positive, n (%) | 10 (50) |
|  | HPV18 positive, n (%) | 2 (10) |
|  | HPV16 or 18 positives, n (%) | 10 (50) |

|  |  |
| --- | --- |
| HR-HPV positives, n (%) | 18 (90) |
| LR-HPV positives, n (%) | 9 (45) |
| Multiple HPV infections | 15 (75) |

SD: standard deviation. CIN: cervical intraepithelial neoplasia. HR-HPV: high-risk HPV (HPV16, 18, 31, 33, 35, 39, 45, 51, 52, 56, 58, 59, and 68). LR-HPV: low-risk HPV (6, 11, 40, 42, 54, 61, 62, 71, 72, 81, 83, 84, and CP 6108)

### DNA yield

DNA yield per 100 μL ThinPrep solution was 0.09 ± 0.06 μg in ZymoBIOMICS, 0.04 ± 0.01 μg in PowerFecalPro, and 0.21 ± 0.23 μg in QIAampMini. DNA yield was not calculated for IndiSpin, as Poly-A Carrier DNA was used. The DNA yield of PowerFecalPro was significantly lower than that of ZymoBIOMICS (adjusted p value < 0.001) and QIAampMini (adjusted p value < 0.001) based on Dunn’s test with Benjamini-Hochberg-adjustment (Figure S1).

### Mock and Negative Controls

We observed that we were able to reasonably recover the expected taxa of our mock community positive control from the American Type Culture Collection (ATCC MSA1007). Each of the following taxa should have a relative abundance of ∼16.7% of the total sample. It should be noted that factors, such as sample preparation and primer biases, can cause deviations from the expected mock community [76] [77] [78]. We observed: 7.184% *Gardnerella spp*., 14.807% *Lactobacillus jensenii* (40.185% *Lactobacillus spp.*), 16.530% *Mycoplasma hominis*, 14.311% *Prevotella bivia* (14.327 % *Prevotella spp.*), and 21.429% *Streptococcus agalactiae* (21.449% *Streptococcus spp*.). A total of 127,193 reads were generated from the mock community control. Of which, 99.68% (126,783 / 127,193 reads) were from expected members of the mock community. For the negative control (ThinPrep preservation solution) only 1,791 reads were generated. 1,400 reads were from *Staphylococcus spp.*, 323 reads were from *Micrococcus spp.*, and 47 reads were *Lactobacillus spp.*. The remaining 21 reads were spurious.

### Number of reads and Operational Taxonomic Units (OTUs) before rarefying

We obtained a total of 11,149,582 reads for 80 DNA extractions. IndiSpin (168,349 ± 57,451 reads) produced a significantly higher number of reads compared to PowerFecalPro (115,610 ± 68,201 reads, p value = 0.020, Dunn’s test with Benjamini-Hochberg-adjustment) as shown in Table 3 Approximately 90% of reads were assigned to gram-positive bacteria and about 10% of reads were assigned to gram-negative bacteria across all kits.

**Table 3.**
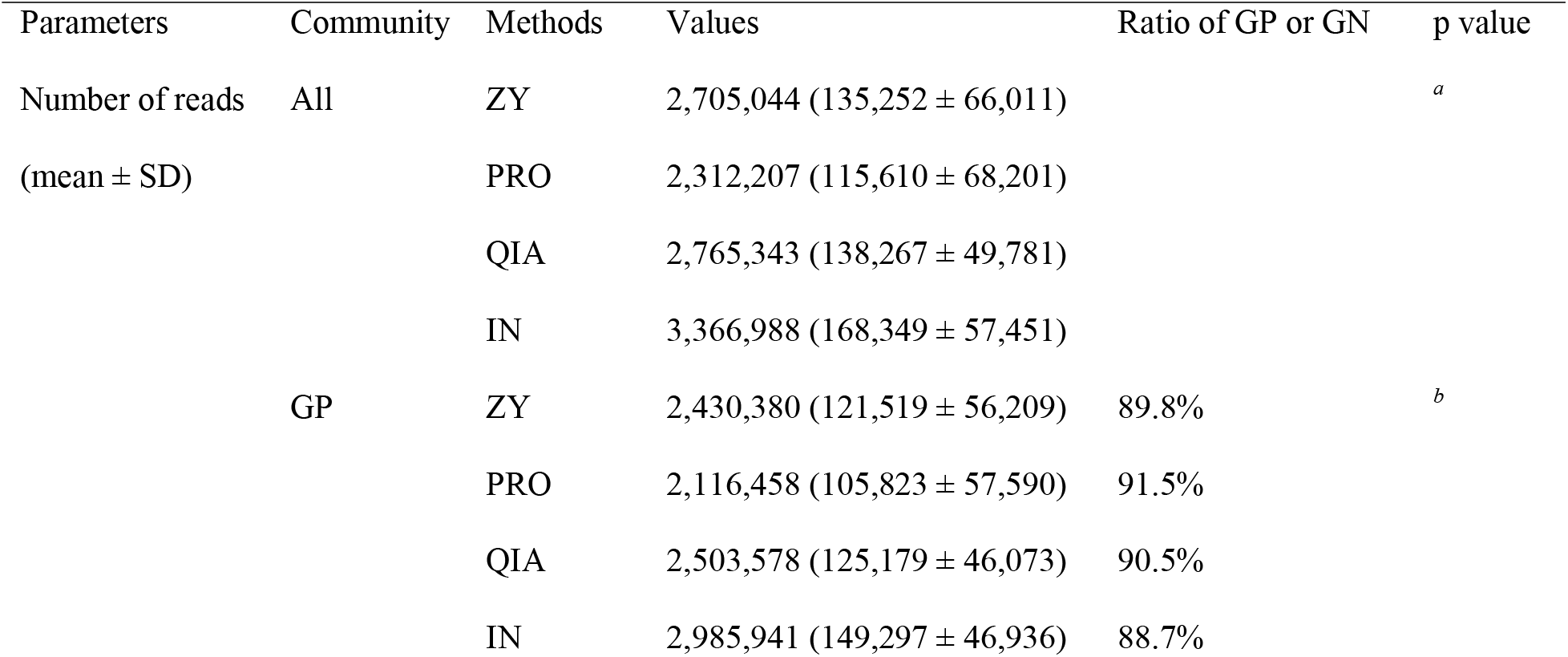
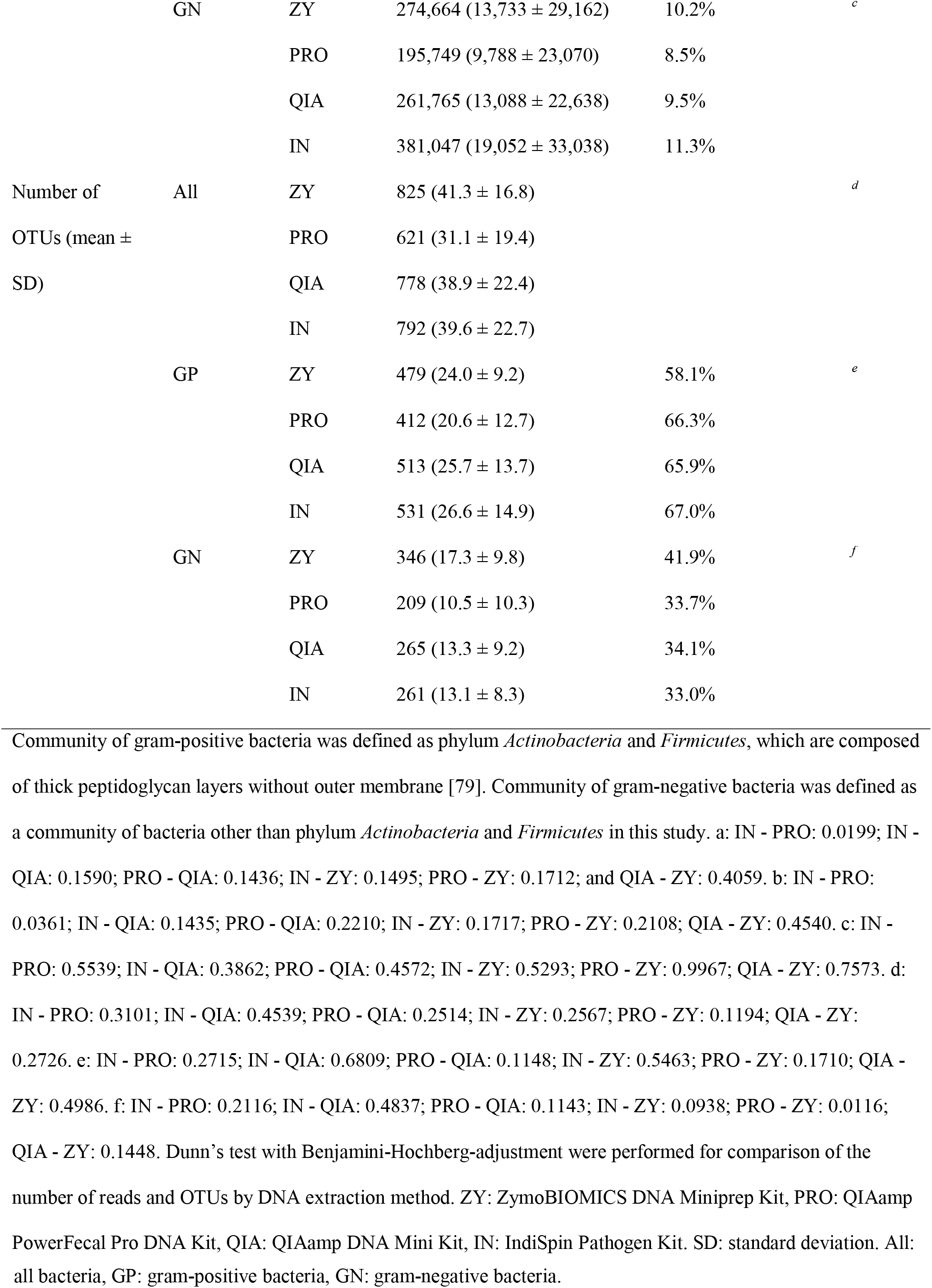
Reads and OTUs before rarefying assigned to all, gram-positive, and gram-negative bacteria per DNA extraction protocols.

Prior to rarefying, the ZymoBIOMICS kit captured a greater representation of gram-negative bacterial OTUs (total 346, 17.3 ± 9.8) compared to PowerFecalPro (total 209, 10.5 ± 10.3, p value = 0.012, Dunn’s test with Benjamini-Hochberg-adjustment, ratio of gram-negative bacteria: 41.9% vs 33.7%) as shown in Table 3. No significant differences in the number of OTUs before rarefying was detected for the entire bacterial community or gram-positive bacteria.

### Microbiome composition per DNA extraction protocol

We analyzed whether differences in DNA extraction methods affect our ability to assess cervical microbiota composition. The patients can be identified by whether or not they displayed a *Lactobacillus*-dominant community type (Figures 2A & S4). Variation between patients was a significantly greater influence on the observed microbial composition than was the method of DNA extraction (F.Model: 199.4, R2: 0.982, and p value: 0.001 for patients vs F.Model: 2.9, R2: 0.003, and p value: 0.002 for DNA extraction, Adonis test, Figure 2A).

**Figure 2.**
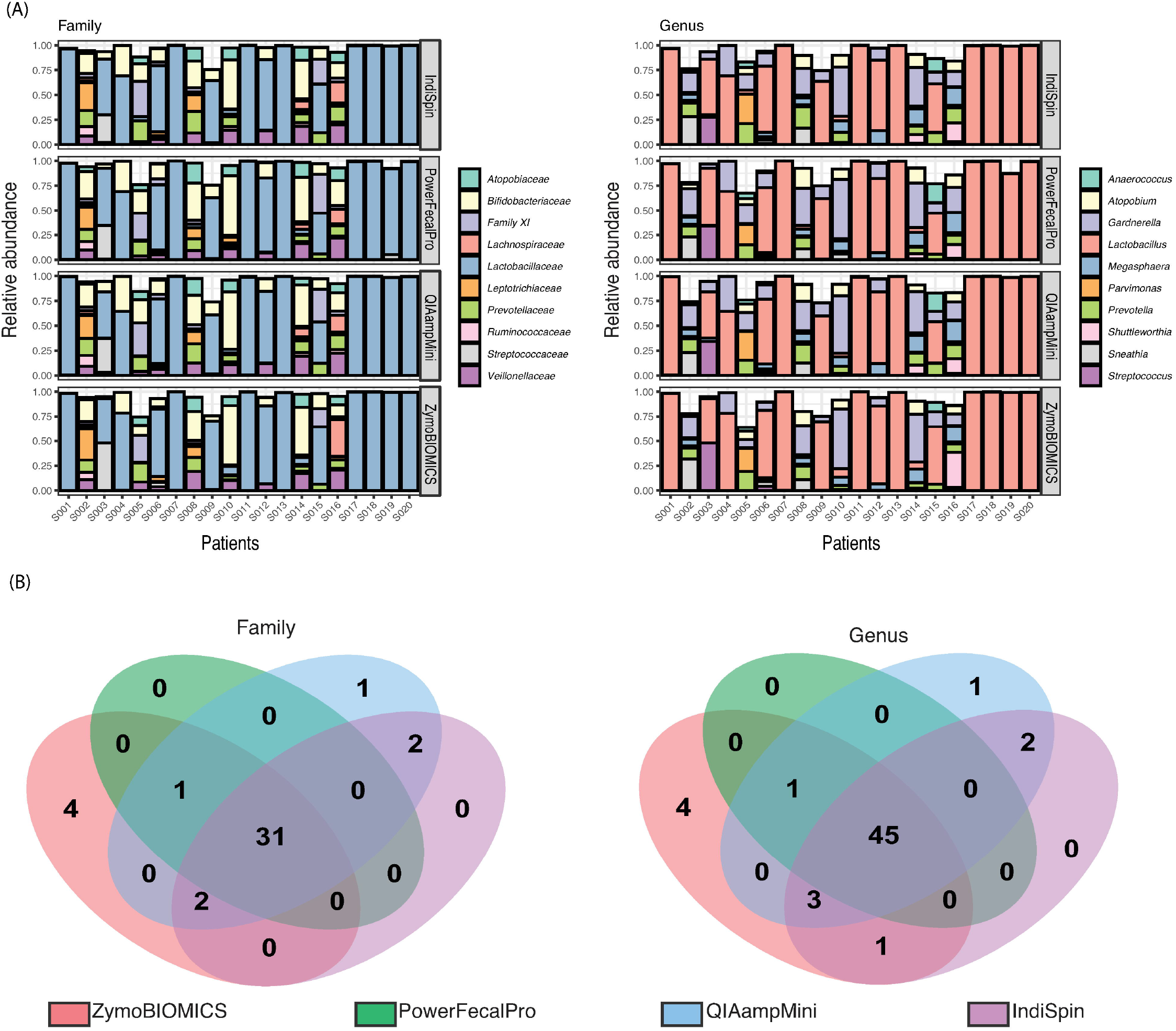
Taxonomic resolution among DNA extraction protocols. (A) Relative abundance of microbe at family level (left) and genus level (right) per DNA extraction method showed the pattern that variance of microbe composition per patient was higher than that per DNA extraction protocol. These patterns were confirmed by values of Adonis test (`q2-diversity adonis`); F.Model: 199.4, R2: 0.982, and p value: 0.001 for patients and F.Model: 2.9, R2: 0.003, and p value: 0.002 for DNA extraction [60] [61]. After all count data of taxonomy were converted to relative abundance as shown in the y-axis, the top ten taxonomy at each family and genus level were plotted in colored bar plot and other relatively few taxonomies were not plotted. The 20 patients ID were described in the x-axis. (B) Venn diagrams, considering only those OTUs with a frequency greater than 0.005% shown, revealed that ZymoBIOMICS had four unique taxa at family (left) and genus (right) taxonomic level. Thirty-one of 41 families and 45 of 57 genera were detected with all DNA extraction protocols.

The following top 10 abundant families are shown in Figure 2A (left) and constituted approximately 95.7% of cervical bacteria in all kits (80 DNA extractions); *Lactobacillaceae* (58.9%), *Bifidobacteriaceae* (13.7%), *Veillonellaceae* (4.8%), *Prevotellaceae* (4.3%), *Family XI* (3.9%), *Atopobiaceae* (3.0%), *Leptotrichiaceae* (2.5%), *Streptococcaceae* (2.0%), *Lachnospiraceae* (1.6%). *Ruminococcaceae* (0.9%). The following top 10 abundant genera are shown in Figure 2A (right) and constituted approximately 92% of cervical bacteria; *Lactobacillus* (58.9%), *Gardnerella* (13.6%), *Prevotella* (4.2%), *Megasphaera* (3.7%), *Atopobium* (3.0%), *Sneathia* (2.5%), *Streptococcus* (1.9%), *Parvimonas* (1.7%), *Shuttleworthia* (1.4%), and *Anaerococcus* (1.1%).

### Shared and unique microbiota among DNA extraction protocols

All DNA extraction methods were generally commensurate with one another, there were 31 of 41 shared microbes at the family level (Figure 2B left) and 45 of 57 shared microbes at the genus level (Figure 2B right) among the DNA extraction protocols.

However, four gram-negative taxa were uniquely detected by ZymoBIOMICS and one taxon was uniquely detected by QIAampMini both at the genus level (Figure 2B right). Of the uniquely detected ZymoBIOMICS OTUs, *Methylobacterium* was detected in 5 of the 80 DNA extractions, consisting of 912 reads: 0.01% of all kit extractions. A member of this genus, *Methylobacterium aerolatum*, has been reported to be more abundant in the endocervix than the vagina of healthy South African women [80]. *Bacteroidetes*, which are often reported as enriched taxa in an HIV positive cervical environment [81], was detected in 12 of the 80 DNA extractions (1,028 reads; 0.01%). *Meiothermus* was detected in 9 of the 80 DNA extractions (882 reads; 0.01%) and *Hydrogenophilus* was detected in 14 of the 80 DNA extractions (2,488 reads, 0.02%). *Meiothermus* and *Hydrogenophilus* [82] are not considered to reside within the human environment, and are likely kit contaminants, as previously reported [83]. A unique gram-positive taxon obtained from the QIAampMini, *Streptomyces*, which was reported to be detected from the cervicovaginal environment in the study of Kenyan women [84], was detected in all 20 of the QIAampMini DNA extractions (6,862 reads; 0.06%). No unique taxa were detected in PowerFecalPro and IndiSpin. Although less than 0.005% of the total data set, two samples of IndiSpin also detected potential kit contaminant, *Tepidiphilus* (*Hydrogenophilaceae*).

### Alpha and beta diversity

The observed alpha diversity was similar across all kits except for a few cases (Figure 3). Significantly higher species richness (`q2-breakaway`) was observed between the ZymoBIOMICS (56.1 ± 19.4) protocol and that of PowerFecalPro (43.2 ± 32.9, p = 0.025) (Figure 3). Similarly, Faith’s Phylogenetic Diversity was observed to be higher with the ZymoBIOMICS protocol (6.6 ± 2.2), compared to PowerFecalPro (4.5 ± 1.9, p = 0.012). The use of IndiSpin also resulted significantly higher alpha diversity than that of PowerFecalPro in an analysis of Species richness (p = 0.042). Non-phylogenetic alpha diversity metrics such as Observed OTUs, Shannon’s diversity index, and Pielou’s Evenness did not show differences among the four methods.

**Figure 3.**
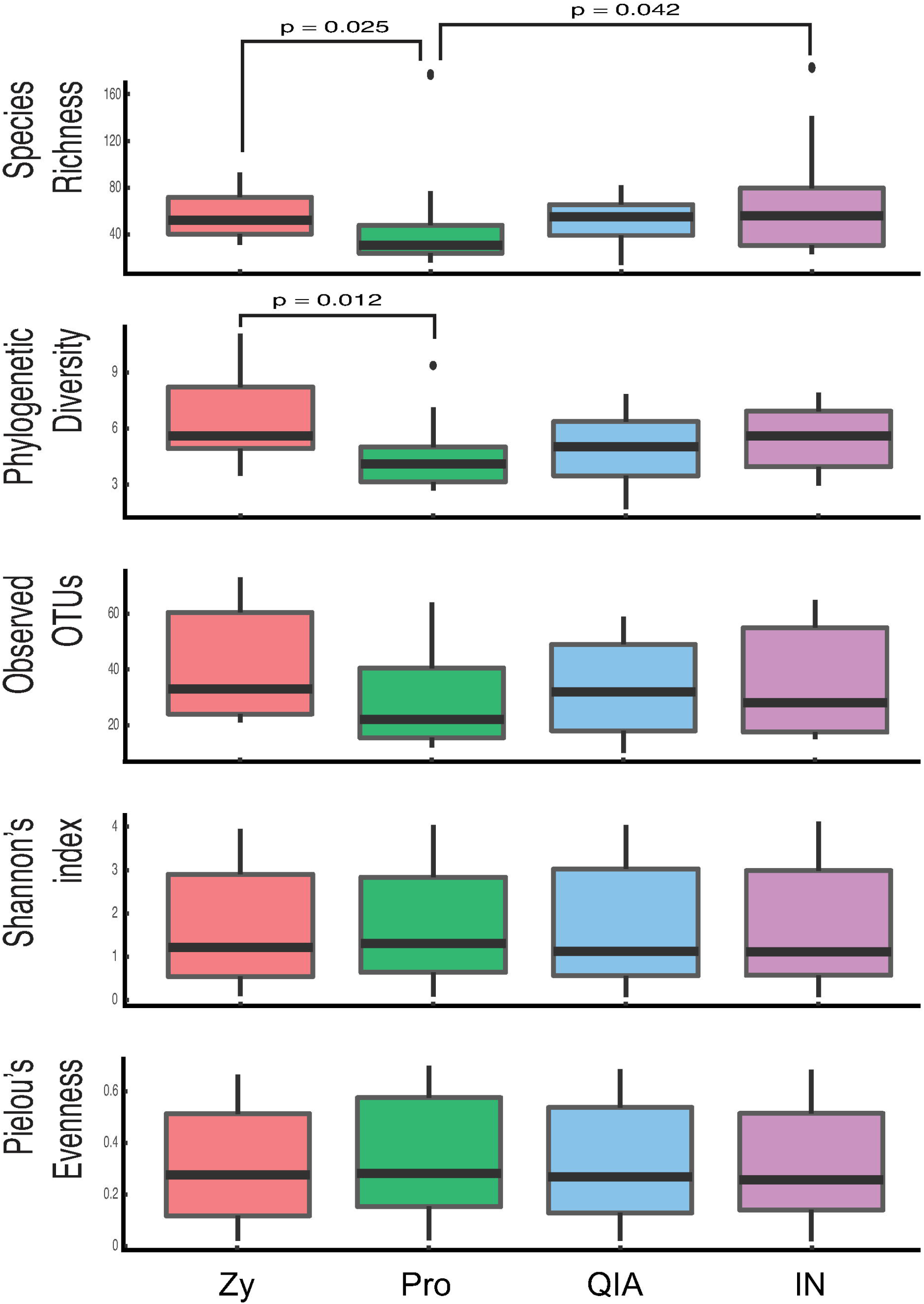
Comparisons of alpha diversity between different DNA extraction protocols. The alpha diversity indices determined by Species richness and Phylogenetic diversity are significantly higher with ZymoBIOMICS in comparison with PowerFecalPro (p = 0.025 and 0.012, respectively, Dunn’s test with Benjamini-Hochberg-adjustment). IndiSpin also showed significantly higher diversity than that of PowerFecalPro using analysis of Species richness (p = 0.042, Dunn’s test with Benjamini-Hochberg-adjustment). No significant differences were observed in other alpha diversity indexes such as observed OTUs, Shannon’s diversity index, and Pielou’s Evenness. Zy: ZymoBIOMICS DNA Miniprep Kit, Pro: QIAamp PowerFecal Pro DNA Kit, QIA: QIAamp DNA Mini Kit, IN: IndiSpin Pathogen Kit.

Similar to the alpha diversity results above, no significant differences were observed with other metrics, including `q2-deicode` (Aichison distances). Only qualitative metrics such as Unweighted UniFrac and Jaccard distance, revealed significant differences in a few cases (Table 4 Figure S2, & Figure S3). Most observed differences were between ZymoBIOMICS and other DNA extraction methods with when qualitative metrics such as Unweighted UniFrac (PowerFecalPro: q = 0.002; QIAampMini: q = 0.002; and IndiSpin: q = 0.002) and Jaccard distances (QIAampMini: q = 0.018 and IndiSpin: q = 0.033) were used. With PowerFecalPro vs. IndiSpin in Unweighted UniFrac (q = 0.023), being the only other observed significant difference. All other metrics showed no significance differences with regard to beta diversity.

**Table 4.**
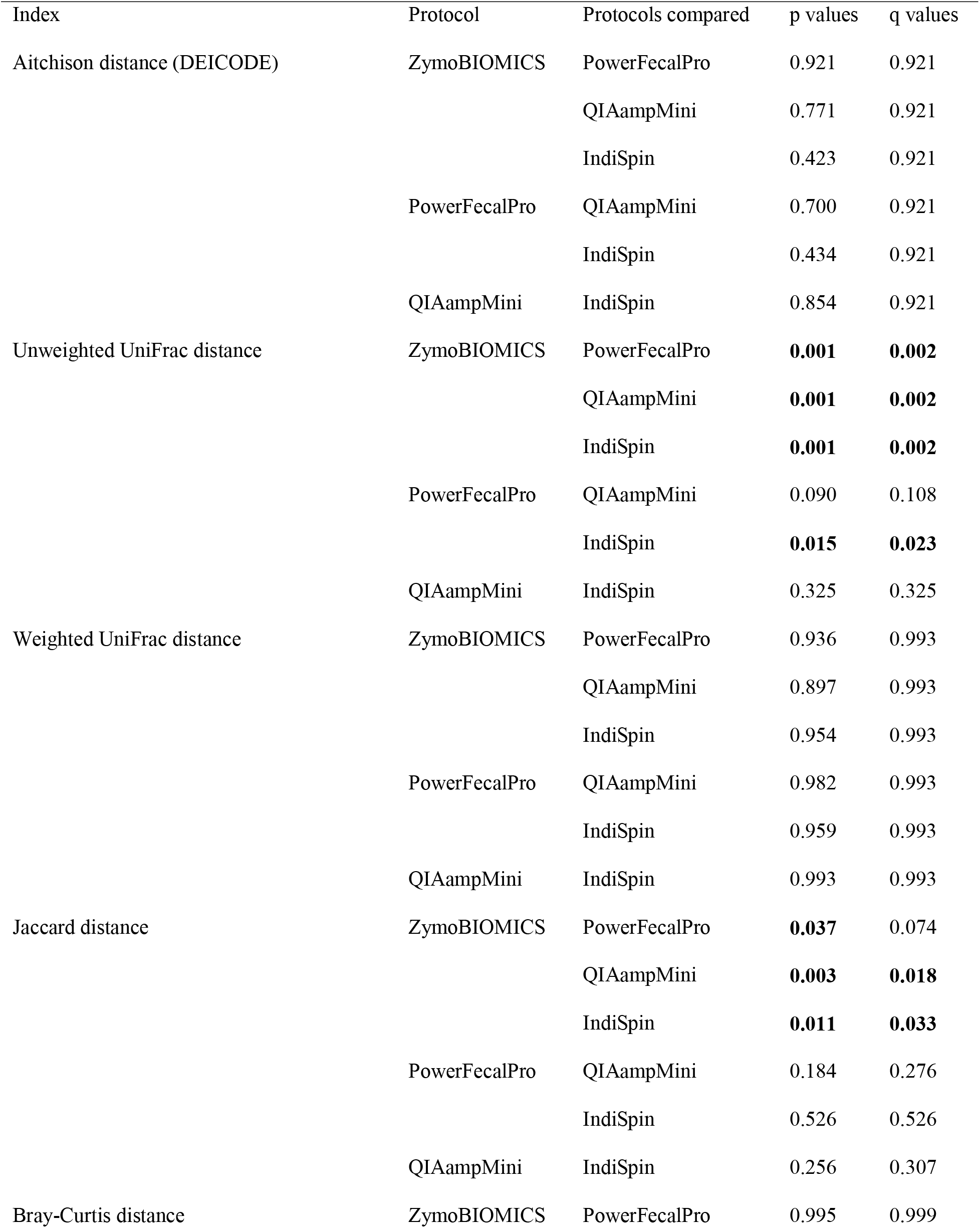
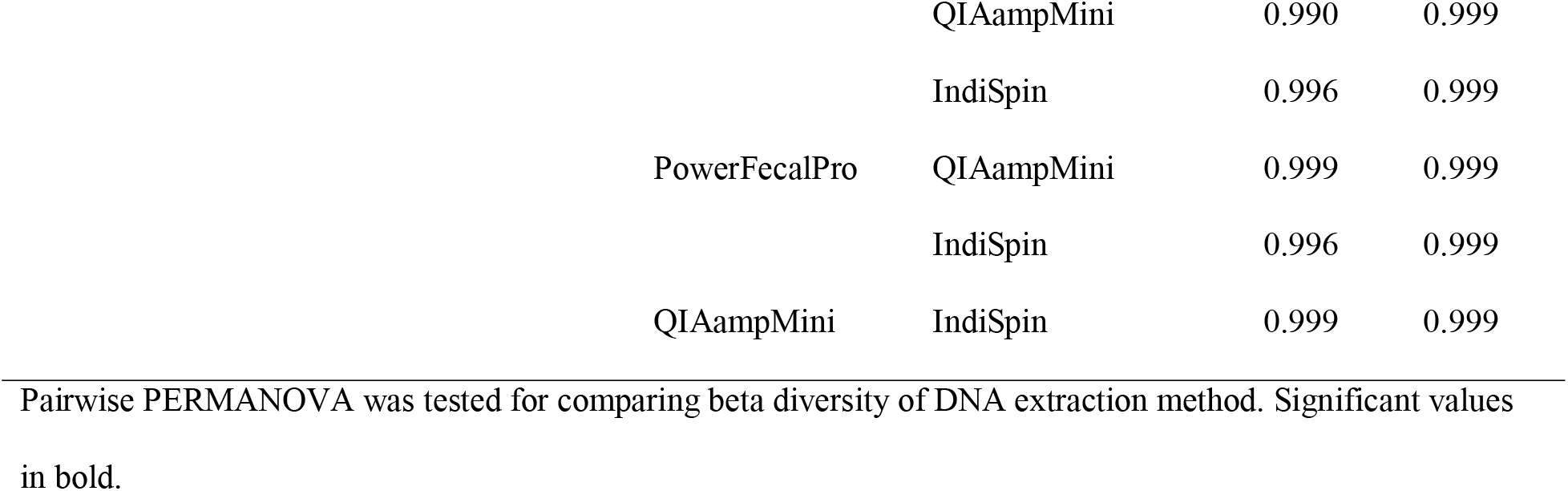
Beta diversity among DNA extraction methods.

| Index | Protocol | Protocols compared | p values | q values |
| --- | --- | --- | --- | --- |
| Aitchison distance (DEICODE) | ZymoBIOMICS | PowerFecalPro | 0.921 | 0.921 |
|  |  | QIAampMini | 0.771 | 0.921 |
|  |  | IndiSpin | 0.423 | 0.921 |
|  | PowerFecalPro | QIAampMini | 0.700 | 0.921 |
|  |  | IndiSpin | 0.434 | 0.921 |
|  | QIAampMini | IndiSpin | 0.854 | 0.921 |
| Unweighted UniFrac distance | ZymoBIOMICS | PowerFecalPro | **0.001** | **0.002** |
|  |  | QIAampMini | **0.001** | **0.002** |
|  |  | IndiSpin | **0.001** | **0.002** |
|  | PowerFecalPro | QIAampMini | 0.090 | 0.108 |
|  |  | IndiSpin | **0.015** | **0.023** |
|  | QIAampMini | IndiSpin | 0.325 | 0.325 |
| Weighted UniFrac distance | ZymoBIOMICS | PowerFecalPro | 0.936 | 0.993 |
|  |  | QIAampMini | 0.897 | 0.993 |
|  |  | IndiSpin | 0.954 | 0.993 |
|  | PowerFecalPro | QIAampMini | 0.982 | 0.993 |
|  |  | IndiSpin | 0.959 | 0.993 |
|  | QIAampMini | IndiSpin | 0.993 | 0.993 |
| Jaccard distance | ZymoBIOMICS | PowerFecalPro | **0.037** | 0.074 |
|  |  | QIAampMini | **0.003** | **0.018** |
|  |  | IndiSpin | **0.011** | **0.033** |
|  | PowerFecalPro | QIAampMini | 0.184 | 0.276 |
|  |  | IndiSpin | 0.526 | 0.526 |
|  | QIAampMini | IndiSpin | 0.256 | 0.307 |
| Bray-Curtis distance | ZymoBIOMICS | PowerFecalPro | 0.995 | 0.999 |

|  |  |  |  |
| --- | --- | --- | --- |
|  | QIAampMini | 0.990 | 0.999 |
|  | IndiSpin | 0.996 | 0.999 |
| PowerFecalPro | QIAampMini | 0.999 | 0.999 |
|  | IndiSpin | 0.996 | 0.999 |
| QIAampMini | IndiSpin | 0.999 | 0.999 |

Pairwise PERMANOVA was tested for comparing beta diversity of DNA extraction method. Significant values in bold.

### Differential accessibility of microbiota by DNA extraction protocol

Linear discriminant analysis (LDA) effect size (LEfSe) analysis [62], identified several taxonomic groups, defined with an LDA score of 2 or higher *(one-against-all*), for differential accessibility by extraction kit: 23 in ZymoBIOMICS, 0 in PowerFecalPro, 3 in QIAampMini, and 3 in IndiSpin (Figure 4A). The following taxa were found to be highly accessible (LDA score > 3) with the use of the ZymoBIOMICS kit: Phylum *Proteobacteria*, Class *Gammaproteobacteria*, Order *Betaproteobacteriales*, Family *Bacillaceae*, and Genus *Anoxybacillus*. Whereas the Order *Streptomycetales* was highly enriched with the use of the QIAampMini (LDA score > 3). However, post-hoc analysis of the LEfSe output, using Scheffe test [64] revealed that only the contaminants *Meiothermus* and *Hydrogenophilus* were enriched with Zymo, and *Streptomyces* was enriched in QIA (Figure 4A). These results reveal minimal to no significant enrichment of specific microbiota across extraction kits.

**Figure 4.**
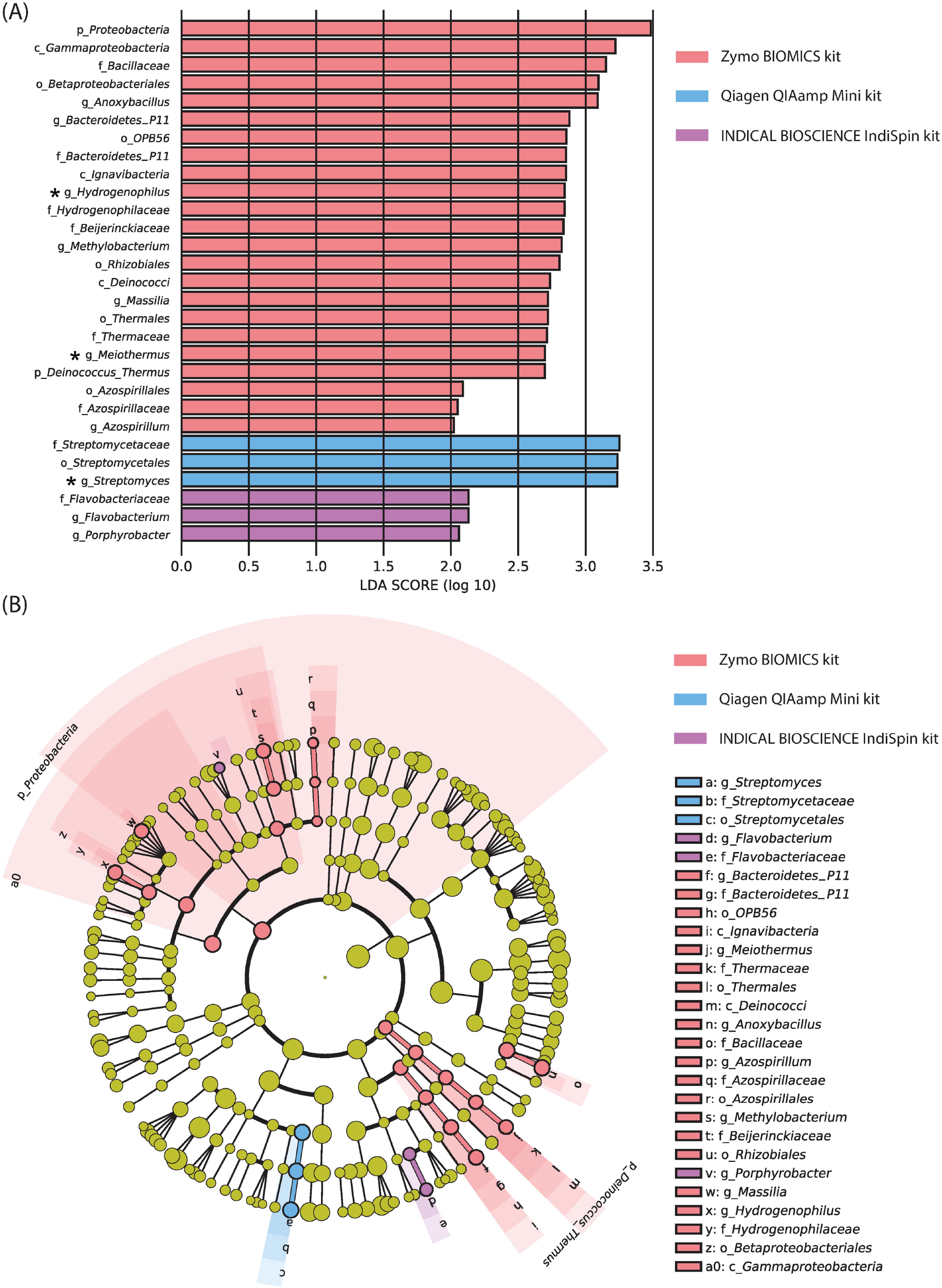
Distinct detections of microbe among the DNA extraction protocols. (A) A bar graph showing 23 significantly enriched taxa with ZymoBIOMICS, 3 with QIAamp DNA Mini Kit, and 3 with IndiSpin Pathogen Kit determined by the linear discriminant analysis (LDA) effect size (LEfSe) analyses [62]. Asterisks denote taxa of genus level that were significant after post-hoc significant testing with Scheffe. (B) A taxonomic cladogram from the same LEfSe analyses showing that the significantly enriched microbiota in ZymoBIOMICS were composed of phylum *Proteobacteria*. Also note that *Meiothermus* (a member of the phylum *Deinococcus-Thermus*) *Hydrogenophilaceae* (a member of the phylum *Proteobacteria*), and *Hydrogenophilus* (a member of the phylum *Proteobacteria*) are likely an extraction kit contaminant. Zy: ZymoBIOMICS DNA Miniprep Kit, Pro: QIAamp PowerFecal Pro DNA Kit, QIA: QIAamp DNA Mini Kit, IN: IndiSpin Pathogen Kit. g_: genus, f_: family, o_: order, c_: class, p_: phylum.

### Microbial community type and HPV16

Dirichlet Multinomial Mixtures (DMM) model [70] detected two cervical microbial community types across all four DNA extraction protocols (Figure S4). Community type I was composed of the following: *Gardnerella sp.* (ZymoBIOMICS: 17.1%; PowerFecalPro: 20%; QIAampMini: 23%; IndiSpin: 20%), *Lactobacillus iners* (ZymoBIOMICS: 6.3%; PowerFecalPro: 5%; QIAampMini: 6%; IndiSpin: 5%), *Atopobium vaginae* [10] (ZymoBIOMICS: 3.5%; PowerFecalPro: 3%; QIAampMini: 4%; IndiSpin: 5%), *Clamydia trachomatis* (ZymoBIOMICS: 1.9%; PowerFecalPro: 2%; QIAampMini: 3%; IndiSpin: 2%), *Shuttleworthia sp.* (ZymoBIOMICS: 1.8%; PowerFecalPro: 2%; QIAampMini: 2%; IndiSpin: 2%). Some members of *Shuttleworthia* are considered to be bacterial vaginosis□associated bacterium (BVAB) [85], further investigation is required to determine if this OTU is indeed a BVAB. We determined this community type “high diversity type”. Community type II was is dominated by *Lactobacillus iners* at 88%, 85%, 83%, and 85% respectively for ZymoBIOMICS, PowerFecalPro, QIAampMini, and IndiSpin.

The relationship between HPV16 infection and community type was observed to be significantly associated with community type I (HPV16 positive patients [n = 9], HPV16 negative patients [n = 1]) and not community type II (HPV16 positive patients [n = 1], HPV16 negative patients [n = 9], p = 0.001, Fisher’s exact test) regardless of the DNA extraction kit used (Figure S4A). In support of this result, analysis of differentially abundant microbiota using `q2-aldex` (Benjamini-Hochberg corrected p value of Wilcoxon test: p < 0.001, standardized distributional effect size: −1.2) revealed that *Lactobacillus iners* were differentially enriched in the cervical environment without HPV16. LEfSe analysis also detected that genus *Lactobacillus* were enriched in the cervical environment without HPV16 (p < 0.001, LDA score: 5.38, Figure S4B). No significant differences were observed in the relationship between community type and HPV18 (p = 0.474, Fisher’s exact test), HR-HPV (p = 0.474, Fisher’s exact test), LR-HPV (p = 0.370, Fisher’s exact test), multiple HPV infections (p = 0.303, Fisher’s exact test), results of cervical biopsy (p = 0.554, Fisher’s exact test), and race (African Americans vs not-African Americans: p = 1; European descent vs non-European descent: p = 0.656; Hispanic vs non-Hispanic: p = 0.350, Fisher’s exact test, Figure S4A).

## Discussion

In this study, we evaluated the utility of LBC specimens for the collection and storage of cervical samples for microbiome surveys based on the 16S rRNA marker gene. We simultaneously compared the efficacy of several commonly used DNA extraction protocols on these samples in an effort to develop a standard operating procedure/protocol (SOP) for such work. We’ve also been able to show that there are two cervical microbial community types, which are associated with the dominance or non-dominance of *Lactobacillis iners* and HPV16 status (Figures 2A & S4A). The relationship between community types and HPV16 was detected regardless of the DNA extraction protocol used.

This study evaluated the composition of microbiota accessible across all DNA extraction methods. All kits were commensurate in their ability to capture the microbial composition of each patient and the two observed cervical microbial community state types, making all of these protocols viable for discovering broad patterns of microbial diversity. It should be noted, however, that a singular kit should be used through the entirety of a study to minimize any subtle differences between samples, particularly when qualitative or richness-based diversity metrics are used. We detected potential DNA contamination with the ZymoBIOMICS and IndiSpin kits. The number of OTUs prior to rarefying revealed that the ZymoBIOMICS protocol detected more gram-negative OTUs than the PowerFecalPro (Figure 2B & Table 3). In particular, LEfSe analysis has shown that phylum *Proteobacteria* can be better detected with the ZymoBIOMICS kit (Figure 4). This signature was no longer observed after *post hoc* testing.

Although rarefying microbiome data can be problematic [86], it can still provide robust and interpretable results for diversity analysis [87], we were able to observe commensurate findings with non-rarefying approaches such as `q2-breakaway` [65], `q2-deicode` [66], and LEfSe [62]. Beta-diversity analysis via Unweighted UniFrac also revealed that ZymoBIOMICS was significantly different from all other kits (Table 4). There were no differences in non-phylogenetic indices of alpha diversity (Figure 3). These findings lead us to surmise that qualitative metrics are more sensitive to differences between extraction kits, while quantitative metrics were more sensitive to differences between subject (Figures S2 & S3).

Although we hypothesized that the detection of difficult-to-lyse-bacteria (*e.g*. gram-positive bacteria) would vary by kit, we observed no significant differences (Table 3). The number of reads of gram-positive and gram-negative bacteria also showed that there was no difference in the four kits (Table 3). This is likely due to several modifications made to the extraction protocol as outlined in Table 1. That is, we added bead beating and mutanolysin to the QIAampMini protocol [36]. We also modified the beating time of the ZymoBIOMICS kit down to 2 minutes from 10 minutes (the latter being recommended by the manufacturer) to minimize DNA shearing. We may use the extracted DNA from ZymoBIOMICS for long-read amplicon sequencing platforms such as PacBio (Pacific Biosciences of California, Inc) [88] or MinION (Oxford Nanopore Technologies) [89] [90]. Excessive shearing can render these samples unusable for long-read sequencing. It is quite possible that we could have observed even more diversity with the ZymoBIOMICS kit for our amplicon survey if we conducted bead-beating for the full 10 minutes.

One limitation of our study is the lack of fresh LBC samples that would have enabled assessment the effects of prolonged storage on determining microbial community composition due to potential DNA degradation [25]. We think this may be unlikely, as our LBC samples were immediately frozen in −80 □C, and DNA degradation within LBC samples stored at −80 □C has been shown to be minimal [26]. However, the possibility that the observed microbial community composition may not be indicative of the community at the time of sampling remains. Despite this, our observations are commensurate with several prior studies in this area as outlined below. Community typing and detection of the differentially abundant microbiota revealed that *Lactobacillus iners* were more abundant in the cervical ecosystem without HPV16 (Figure S4). These findings are congruent with those of, Usyk *et al*. [91], Lee *et al*. [1], and Audirac-Chalifour *et al*. [92]. Usyk *et al*., reported that *L. iners* was associated with clearance of HR-HPV infections [91]. Lee *et al*. reported that *L. iners* were decreased in HPV positive women [1]. Also, the results indicated that the proportion of *L. iners* was higher in HPV-negative women compared to HPV-positive women (relative abundance 14.9% vs 2.1%) was reported by Audirac-Chalifour *et al* [92]. Similarly, Tuominen *et al*. [20] reported that *L. iners* were enriched in HPV negative samples (relative abundance: 47.7%) compared to HPV positive samples (relative abundance: 18.6%, p value = 0.07) in the study of HPV positive-pregnant women (HPV16 positive rate: 15%) [93]. As established by the seminal study of Ranjeva *et al*. [94], a statistical model revealed that colonization of specific HPV types including multi-HPV type infection depends on host-risk factors such as sexual behavior, race and ethnicity, and smoking. It is unclear whether the association between the cervical microbiome, host-specific traits, and persistent infection of specific HPV types, such as HPV16, can be generalized and requires further investigation.

We focused on LBC samples as this is the recommended method of storage for cervical cytology [95]. We used a sample volume of 200 or 300 μL ThinPrep solution in this study. The Linear Array HPV Genotyping Test (Roche Diagnostics) stably detects β-globin with a base length of 268 bp as a positive control. Therefore, using a similar sample volume as HPV genotyping (250 μL), it was expected that V4 (250 bp), which is near the base length of β-globin, would be PCR amplified. It has been pointed out by Ling *et al*. [96] that the cervical environment is of low microbial biomass. To control reagent DNA contamination and estimate the sample volume, DNA quantification by qPCR before sequencing is recommended [97]. Mitra *et al* determined a sample volume of 500 μL for ThinPrep by qPCR in the cervical microbiome study comparing sampling methods using cytobrush or swab [21]. The average storage period from sample collection via LBC to DNA extraction was about two years in this study. Kim *et al*. reported that DNA from the cervix stored in ThinPrep at room temperature or −80°C was stable for at least one year [26]. Meanwhile, Castle *et al*. reported that β-globin DNA fragments of 268 bases or more were detected by PCR in 90 % (27 of 30 samples) of ThinPrep samples stored for eight years at an uncontrolled ambient temperature followed by a controlled ambient environment (10–26.7°C) [25]. Low-temperature storage may allow the analysis of the short DNA fragments of the V4 region after even long-term storage, although further research is needed to confirm the optimal storage period in cervical microbiome studies using ThinPrep. SurePath LBC specimens are as widely used as ThinPrep, but the presence of formaldehyde within the SurePath preservation solution raises concerns about accessing enough DNA for analysis as compared to ThinPrep, which contains methanol [98] [99]. It should also be noted that other storage solutions, *i.e*., those using guanidine thiocyanate have been reported for microbiome surveys of the cervix [100] and feces [101]. A weakness of the current study is that we did not examine the reproducibility of our results as each sample was extracted using each kit once as samples were limited in quantity, and we lacked fresh sample controls to assess the effects of prolonged storage to alter microbial community composition. Although several studies, have shown general stability and accessibility of DNA [26] [102] [103], there is potential for DNA degradation for samples not stored at low temperatures [25] [26]. However, the use of actual patient samples rather than mock samples is a strength of our approach.

## Conclusions

In conclusion, regardless of the extraction protocol used, all kits provided equivalent broad accessibility to the cervical microbiome. Observed differences in microbial composition were due to the significant influence of the individual patient and not the extraction protocol. We have shown that the ability to characterize cervical microbiota from LBC specimens is possible, we were limited in our ability to directly assess if the observed microbial community composition would reflect that of a fresh sample. Despite this limitation, we were able to assess the relationship between HPV and the cervical microbiome, also supported by Kim *et al*. [26] and Castle *et al* [25]. Cervical microbiome in patients with HPV16 or HPV18 which causes 70% of cervical cancers and CIN [104] warrants critical future study. Selection and characterization of appropriate DNA extraction methods are important for providing an accurate census of cervical microbiota and the human microbiome in general [27] [28] [29] [36] [26] [25]. Although we found all four extraction kits to be commensurate in their ability to broadly characterize the CM, one singular kit should be used throughout the entirety of a given study. This study lends support to the view that the selection of a DNA extraction kit depends on the questions asked of the data, and should be taken into account for any cervicovaginal microbiome and HPV research that leverages LBC specimens for use in clinical practice [17] [105].

## Acknowledgements

We thank Togo Picture Gallery [39] for stock images shown in Figure 1.

## Authors’ contributions

M.N. designed and supervised this project. Ta.S. and M.S.R. conducted bioinformatics analysis and wrote paper. Ta.S., H.C., and M.N. created the protocol of DNA extraction. M.N., H.C., S.O., W.G., and To.S. provided important feedback. Samples in the clinical trial were collected by W.G. and his associates. DNA extraction was conducted by Ta.S. Sequencing of 16S RNA gene was conducted by S.O.

## Supporting information

**Figure S1. Comparison of DNA yields by DNA extraction protocols.** DNA yield of QIAampMini was significantly higher than that of PowerFecalPro (p < 0.001, Dunn’s test with Benjamini-Hochberg-adjustment). Also, the DNA yield of ZymoBIOMICS was significantly higher than that of PowerFecalPro (p < 0.001, Dunn’s test with Benjamini-Hochberg-adjustment). The amount of DNA was calculated based on the absorbance of nucleic acids measured by Nanodrop One. By the protocol recommended by the manufacturer, nucleic acid (Poly-A carrier) was used in IndiSpin. Therefore, IndiSpin was excluded from the analysis of DNA yield. The amount of DNA yield per 100 μL ThinPrep sample volume were compared. The bar graph shows the mean and standard deviation. Zy: ZymoBIOMICS DNA Miniprep Kit, Pro: QIAamp PowerFecal Pro DNA Kit, QIA: QIAamp DNA Mini Kit.

**Figure S2. Phylogenetic beta-diversity.** Weighted UniFrac (A & B) and Unweighted UniFrac (C & D), PCoA colored by subject ID (top row) and DNA extraction kit (bottom row). Weighted UniFrac clusters samples by subject whereas Unweighted UniFrac appears more sensitive to the type of DNA extraction kit. Data were rarefied to 51,197 reads per sample.

**Figure S3. Deicode (Robust Aitchison PCA) beta-diversity.** Non-rarefaction-based analysis of beta-diversity. Samples are colored by individual subject ID (A) and DNA extraction kit (B). Samples predominately cluster by subject and not DNA extraction kit.

**Figure S4. Community type and HPV 16 assessed by using 4 kits** (A) Community types were classified into two types in all DNA extraction kits, mainly based on the percentage of *Lactobacillus iners*. HPV16 infection was negatively associated with the dominance of *L. iners* (community type I; p = 0.001, Fisher’s exact test) regardless of DNA extraction method. Although, we observed slight variation in the abundance of microbiota across the extraction kits (even within the same individual patient), the ability to detect two community types was identical across all DNA extraction kits. No significant differences were observed in the relationship of other phenotypes of patients (HPV18, HR-HPV, LR-HPV, multiple HPV infections, Biopsy, and Race). The top 15 bacteria detected for each DNA extraction kit are shown. Samples were clustered by the Dirichlet component. Narrow columns show each sample and a broader column shows averages of samples. Rows show taxa at the species level. Dark or thin colors correspond to larger or smaller counts of OTUs, respectively. CT: community type. (B) LEfSe analysis, using combined data from all four kits detected a significant enrichment of 66 taxa in the cervical environment with HPV16 infection and 17 taxa without HPV16 infection. Genus *Lactobacillus* were enriched in the HPV16 negative patients (p < 0.001, LDA score: 5.38). Asterisks denote taxa that were significant after post-hoc significant testing with Scheffe test [64].

