## Supplementary figures and images for "Evaluation of DNA extraction protocols from liquid-based cytology specimens for studying cervical microbiota"

### Figure S1

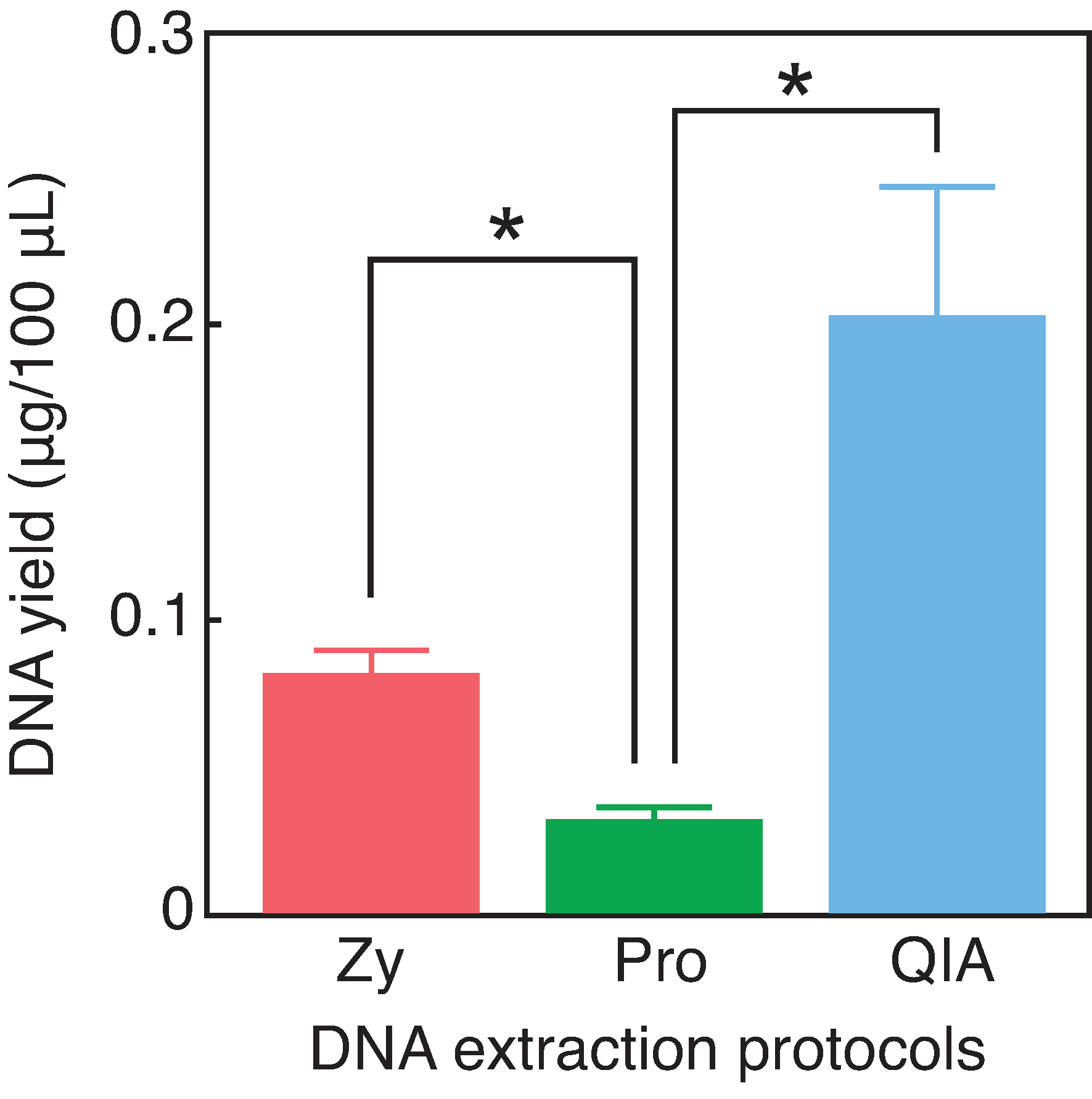

### Figure S2

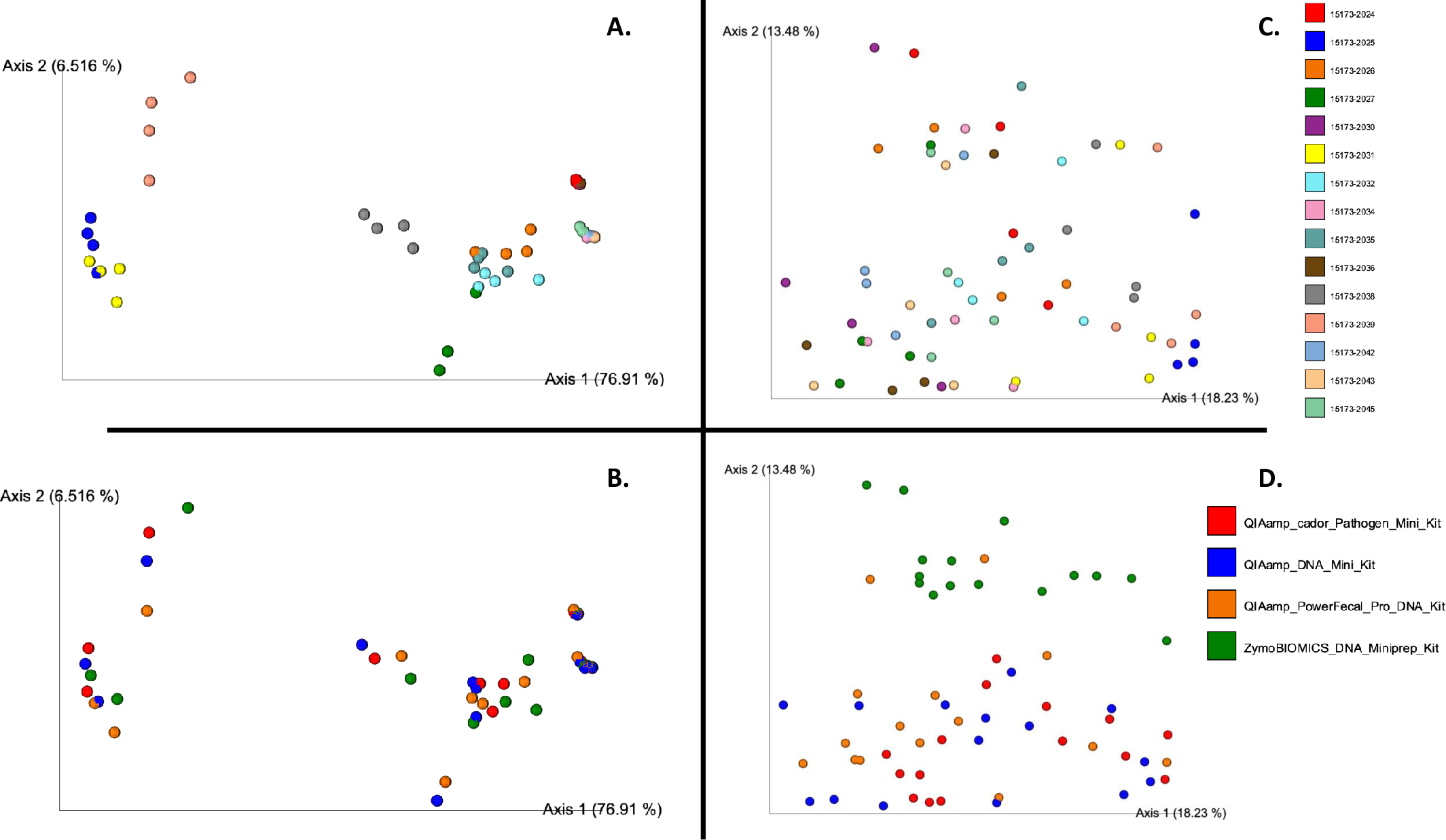

### Figure S3

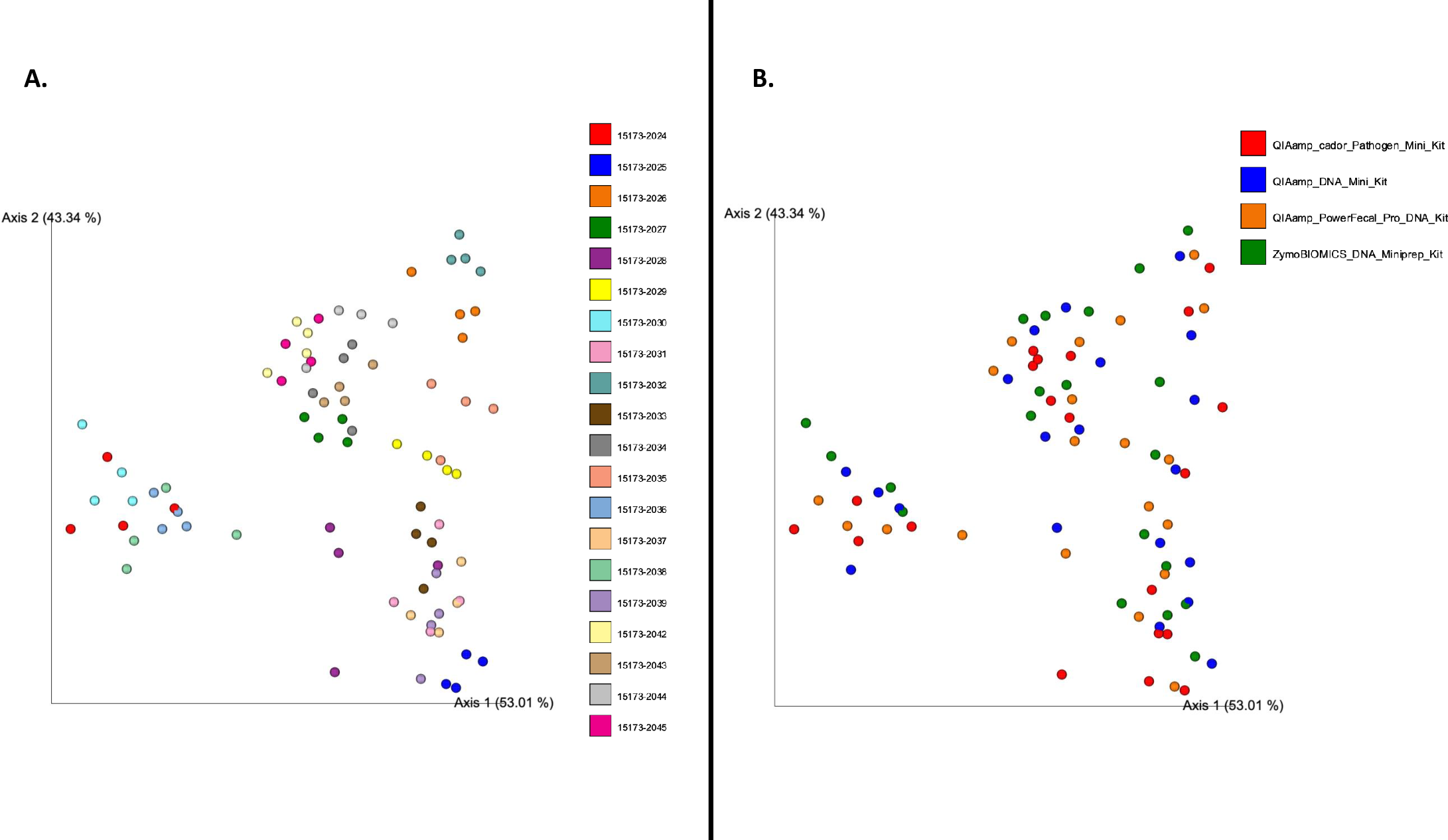

### Figure S4

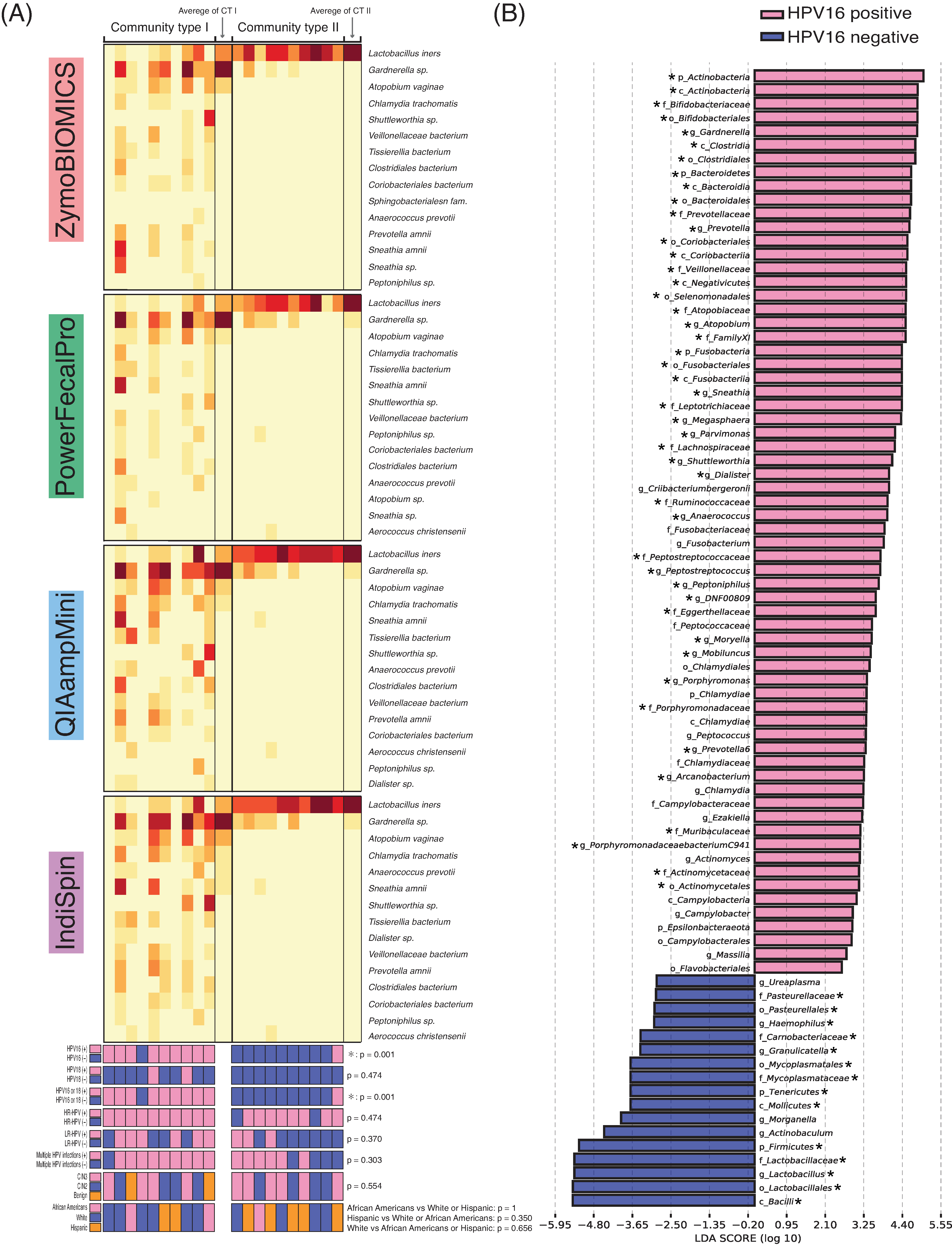
