## Supplementary material for "Evaluation of DNA extraction protocols from liquid-based cytology specimens for studying cervical microbiota": Table S1

| **Table S1. Diversity metrics used in this study.** | | | | | |
| --- | --- | --- | --- | --- | --- |
| **No.** | **Parameter** | **Alpha or Beta diversity** | **Used data with/without rarefying** | **Input data with/without phylogenetic information** | **Plugin of QIIME 2** |
| 1 | Species richness | Alpha | Not rarefied | Non-phylogenetic | q2-breakaway [65] |
| 2 | Faith’s Phylogenetic Diversity | Alpha | Rarefied | Phylogenetic | q2-diversity |
| 3 | Observed OTUs | Alpha | Rarefied | Non-phylogenetic | q2-diversity |
| 4 | Shannon’s diversity index | Alpha | Rarefied | Non-phylogenetic | q2-diversity |
| 5 | Pielou’s Evenness | Alpha | Rarefied | Non-phylogenetic | q2-diversity |
| 6 | Aitchison distance | Beta | Not rarefied | Non-phylogenetic | q2-deicode [66] |
| 7 | Unweighted UniFrac distance | Beta | Rarefied | Phylogenetic | q2-diversity |
| 8 | Weighted UniFrac distance | Beta | Rarefied | Phylogenetic | q2-diversity |
| 9 | Jaccard distance | Beta | Rarefied | Non-phylogenetic | q2-diversity |
| 10 | Bray-Curtis distances | Beta | Rarefied | Non-phylogenetic | q2-diversity |
| 11 | Adonis | Beta | Rarefied | Non-phylogenetic | q2-diversity adonis [60] [61] |
